# Hidden Markov Modeling Reveals Altered Functional Network Dynamics in Multiple Sclerosis

**DOI:** 10.64898/2026.10.15.759440

**Authors:** Khushi C. Hiremath, Diana A. Hobbs, Danielle M. R. Hettish, Elias Khan, Basil P. Alias, Milo M. Lin, Peter T. Fox, Florence L. Chiang

## Abstract

**Purpose:** Multiple sclerosis (MS) is a neurodegenerative and demyelinating disease of the central nervous system. Recent evidence suggests that localized gray matter atrophy in MS may be involved in distributed functional brain networks. However, the dynamic trajectory of functional network alterations in MS remains unclear. The current study extends prior network-mediated findings and aims to investigate the dynamic functional connectivity (dFC) and energetic landscape across an atrophy-prone network in MS.

**Methods:** Twenty MS participants and twenty healthy controls (HC) underwent resting-state fMRI. The atrophy-based functional network (AFN) model was implemented, and time series from atrophy-prone regions were fit to a 12-state hidden Markov model. Each functional network state, a recurring pattern of activity across the AFN, was characterized by the AFN regions with the strongest activation. The states were then grouped into two metastates. Dynamic functional network metrics were compared between groups by permutation testing and correlated with disability and cognition. The energetic landscape of the network was mapped with a pairwise maximum-entropy model. Temporal irreversibility was also measured, which defines the arrow-of-time asymmetry of network dynamics.

**Results:** People with MS showed lower occupancy of state 1, which peaked on the right middle frontal and anterior cingulate cortex. Occupancy and mean lifetime were higher in states 3 and 10 (p < 0.05). In state 3, cortical activity was below the network average, and state 10 peaked on the postcentral gyrus. At the metastate level, MS participants spent more time in metastate M1 and less in M2 (p = 0.04). Within the MS group, a longer interval between visits to state 3 was associated with greater disability (r = 0.57), and a higher switching rate in state 10 with poorer verbal fluency (r = −0.50). The energetic landscape was similar between groups. Temporal irreversibility was lower in MS, mainly across subcortical connections.

**Conclusion:** MS participants dwelled longer in specific dFC states of the AFN, while the rate of transition between states did not differ from healthy controls. Dynamic functional network metrics were associated with disability and cognitive performance and may serve as functional correlates of clinical impairment. The states were similar between groups, whereas the directed drive between states was lower across subcortical connections. This pattern suggests altered network dynamics rather than altered network architecture in MS. Findings from this study characterize the functional network dynamics of neurodegeneration in MS, which may inform future development of network-based approaches for monitoring disease progression.

## 1 INTRODUCTION

Multiple sclerosis (MS) is a neurodegenerative and demyelinating disease of the central nervous system that affects an estimated 2.8 million people worldwide (Walton et al., 2020). MS can lead to physical disability and cognitive impairment, and the rate of disease progression may be variable (Chiaravalloti & DeLuca, 2008). White matter lesions in MS can be detected with high sensitivity using clinical standard-of-care brain MRI, but lesion burden has limited correlation with clinical disability (Barkhof, 2002). Thus, imaging measures that are more closely associated with clinical status may improve disease monitoring in MS.

Growing evidence suggests that disease worsening in MS may be associated with disruption of large-scale functional brain networks (Mahmoudi et al., 2025; Tahedl et al., 2018). One framework, the network degeneration hypothesis (NDH), posits that regions selectively vulnerable to atrophy serve as nodes in functional networks (Seeley et al., 2009). The NDH has been tested in MS and in other neurodegenerative disorders in which network organization was found to be associated with disease severity (Bassett & Sporns, 2017; Chiang et al., 2021). More recently, the NDH was implemented in MS via the atrophy-based functional network (AFN) model (Chiang et al., 2019, 2021, 2023). The AFN model defines atrophy-prone gray matter hub regions in MS and characterizes inter-regional functional covariance (Chiang et al., 2019). The AFN model was previously implemented in resting-state functional magnetic resonance imaging (rsfMRI) as part of a targeted imaging biomarker development strategy in MS (Chiang et al., 2023). In this model, atrophy-prone gray matter was found to exhibit clinically relevant alterations in static functional connectivity (Chiang et al., 2021).

Static functional connectivity has been widely applied in MS and involves averaging the temporal signal across the scan session. However, static functional connectivity does not describe the temporal structure and dynamic behavior of a functional network. Thus, further evaluation of temporal variation that may occur in functional connectivity may provide additional insights into relevant disease effects (Allen et al., 2014; Hutchison et al., 2013). In MS, decreased variation in dynamic functional connectivity has been associated with cognitive impairment (d’Ambrosio et al., 2020). Other dynamic measures, including dwell time, the time spent in a connectivity state, and switching rate, the frequency of transitions between states, have also been linked to physical disability and cognitive decline (Broeders et al., 2022; Romanello et al., 2022). Altered state dynamics have also been reported in MS (Van Schependom et al., 2019; von Schwanenflug et al., 2023) and in other neurological and psychiatric disorders (Wang et al., 2020; Xiao et al., 2025).

Most studies of dynamic functional connectivity in MS have estimated connectivity within sliding windows of fixed length. The results of this approach depend on the choice of window length, because short windows yield unreliable connectivity estimates and long windows can potentially average over brief network states (Hutchison et al., 2013). The hidden Markov model (HMM) overcomes this limitation by modeling the signal as a sequence of recurring states and inferring the most probable state at each time point (Vidaurre et al., 2017). Dynamic functional network metrics derived from the HMM are obtained from a single resting-state acquisition and may serve as candidate markers for disease monitoring in MS.

In the present study, we characterized functional network dynamics in MS to inform disease monitoring. Specifically, we tested whether dynamic functional connectivity of the AFN is altered in MS. We hypothesized that functional network dynamics would be altered in MS relative to healthy controls, and that the degree of alteration would relate to physical disability and cognitive impairment. Investigation of network dynamics to better characterize connectomics in MS may provide further insights into network-based neurodegeneration in MS.

## 2 MATERIALS AND METHODS

### 2.1 Study Participants

Twenty people with relapsing-remitting MS and 20 age and sex-matched healthy controls (HC) were included. The study was approved by the institutional review board, was compliant with the Health Insurance Portability and Accountability Act, and all participants provided written informed consent. Inclusion criteria for participants with MS were an age of 18-50 years and fulfillment of the 2017 revised McDonald criteria for MS. Exclusion criteria were clinical relapse or intravenous steroid use within the preceding month, other structural brain disease, previous brain surgery, excessive imaging artifact, uncontrolled psychiatric illness, claustrophobia, and other contraindications to MRI.

### 2.2 Image acquisition

All participants underwent whole-brain MRI on a 3T system (Magnetom Tim Trio; Siemens Medical Solutions) with a 12-channel head coil for signal reception and the integrated body coil for transmission, using the acquisition protocol described previously (Chiang et al., 2021). Resting-state T2*-weighted blood oxygen level-dependent (BOLD) fMRI was acquired with a multiband gradient-echo echo-planar imaging sequence (repetition time/echo time = 1400/30 ms; flip angle = 52°; field of view = 211 × 211 mm; matrix = 88 × 88; multiband acceleration factor = 3; isotropic voxel size = 2.4 mm), yielding 700 volumes. Participants were instructed to remain awake with their eyes closed and to let their minds wander, and dummy scans were acquired to establish steady-state magnetization. A gradient-echo field map was acquired with matched shimming and acquisition matrix to correct geometric distortion. For anatomic reference and region-of-interest (ROI) definition, a 3D T1-weighted magnetization-prepared rapid gradient-echo image was obtained (repetition time/echo time/inversion time = 1900/2.26/900 ms; flip angle = 9°; field of view = 256 × 256 mm; matrix = 256 × 256; isotropic voxel size = 1 mm).

### 2.3 Image preprocessing

Functional and anatomic images were preprocessed with fMRIPrep version 25.1.4 (Esteban et al., 2019). For each participant, the T1-weighted image was corrected for intensity nonuniformity, skull-stripped, and segmented into gray matter, white matter, and cerebrospinal fluid (CSF). The existing FreeSurfer (version 7.4.1) surface reconstruction was used for anatomic reference. The resting-state series was corrected for head motion, corrected for susceptibility-related distortion using the field map, coregistered to the T1-weighted image, and kept in native acquisition space; the mean CSF and white matter signals were extracted for use as nuisance regressors. To suppress motion-related artifact, independent component analysis-based automatic removal of motion artifacts (Pruim et al., 2015) was applied with fMRIPost-AROMA, and components classified as noise were removed from the native-space series non-aggressively with FSL ((Jenkinson et al., 2012); version 6.0.7). The mean CSF and white matter time series were then regressed out and the temporal mean restored, yielding the denoised native-space rsfMRI data used for ROI time-series extraction.

### 2.4 Implementation of the atrophy-based functional network

The atrophy-based functional network was originally defined by coordinate-based meta-analysis, with each node represented as a sphere centered on a peak atrophy coordinate (Chiang et al., 2021). The present study extends the network to a dynamic analysis. Static connectivity largely averages out brief fluctuations in the signal, but in a dynamic analysis these fluctuations can appear as temporary changes in connectivity (Hutchison et al., 2013). In the cortex, a sphere may extend across neighboring regions, so its mean signal may include fluctuations from tissue outside the intended region. Each node was therefore defined anatomically by segmentation of each participant’s T1-weighted image, so that each node is bounded by the corresponding anatomical region. This yielded 18 ROIs, and FreeSurfer version 7.4.1 was used for segmentation (Fischl, 2012). Cortical ROIs were taken from the Desikan-Killiany-Tourville atlas (Klein & Tourville, 2012) and comprised the insula, postcentral gyrus, and precentral gyrus bilaterally, together with the left superior temporal gyrus and posterior cingulate cortex, the right anterior cingulate cortex (union of the caudal and rostral anterior cingulate labels), and the right middle frontal gyrus (union of the caudal and rostral middle frontal labels). The caudate nucleus and putamen were obtained bilaterally from the FreeSurfer subcortical segmentation (Fischl et al., 2002). Three thalamic ROIs (left and right pulvinar and right mediodorsal nucleus) were derived from the FreeSurfer probabilistic thalamic nuclei atlas (Iglesias et al., 2018): the pulvinar by combining the anterior, medial, lateral, and inferior subnuclei and the mediodorsal nucleus by combining the medial and lateral subnuclei. The left claustrum was obtained with a contrast-agnostic deep-learning segmentation (Mauri et al., 2025). Each segmentation was converted to NIfTI and resampled from FreeSurfer surface space into the participant’s native functional space in a single interpolation step with ANTs ((Avants et al., 2008); nearest-neighbor interpolation), by concatenating the fMRIPrep-derived FreeSurfer-to-T1-weighted transform with the inverse of the BOLD-to-T1-weighted coregistration transform. The reference grid was the participant’s denoised BOLD volume at 2.4 mm isotropic resolution, and each label was then binarized with FSL (Jenkinson et al., 2012).

### 2.5 Time-series extraction

For each participant, the mean BOLD time series of each of the 18 ROIs was obtained by averaging across all voxels within the corresponding binarized native-space mask with FSL. Because each mask is a cortical or subcortical gray matter label from the participant’s own segmentation, sampling was confined to gray matter without applying a separate tissue mask. This yielded an 18 × 700 region-by-time matrix per participant, which served as input to the HMM.

### 2.6 Hidden Markov model

Dynamic functional connectivity was modeled with an HMM implemented in the osl-dynamics toolbox (Gohil et al., 2023). The model represents the BOLD signal as a sequence of discrete, recurring brain states, each a multivariate Gaussian with a state-specific mean activity vector and full 18 × 18 covariance matrix, with transitions governed by a first-order Markov process. Before fitting, the time series of all 40 participants were temporally concatenated, rotated onto their 18 principal components with no components discarded, and standardized to zero mean and unit variance. This preparation is recommended in osl-dynamics for anatomically parcellated fMRI so that the model captures state switching rather than assigning a scan to a single state (Ahrends et al., 2022; Gohil et al., 2023). The number of states was selected by a grid search that evaluated model free energy in a held-out validation sample (80%/20% participant split). Free energy decreased from 6 to 10 states, and models with 10 to 20 states differed from the 12-state model by 0.01 or less (Figure S1). A 12-state model was therefore retained for consistency with prior resting-state fMRI work (Vidaurre et al., 2017). Three of the 12 states were occupied for less than 1% of time points, which indicates that additional states were not required. State posterior probabilities were inferred per participant with the forward-backward algorithm, and a discrete state sequence was obtained by assigning each time point to the most probable state (Figure S3). Subject-specific spatial descriptions of each state were then obtained by re-estimating the state means and covariances per participant by dual estimation.

### 2.7 State characterization and dynamic functional network metrics

Each state was characterized by the mean activation across the 18 ROIs and by the one or two regions with the strongest positive activation. Mean activation was given by the participant-averaged HMM state mean vector after projection back from the principal component space in which the model was fitted. Four dynamic functional network metrics were computed for each participant and state: fractional occupancy, the proportion of time spent in the state; mean lifetime (dwell time), the mean duration of a visit (in seconds); mean interval, the mean time between consecutive visits (in seconds); and switching rate, the number of entries into the state per second (Vidaurre et al., 2017).

### 2.8 Metastate analysis

To assess higher-order temporal organization, metastates were identified from the structure of state transitions (Vidaurre et al., 2017). The group-averaged transition probability matrix among the active states was treated as a weighted, undirected graph, and the states were clustered with the Louvain algorithm (Blondel et al., 2008), which partitioned the states into two metastates (k = 2). The stability of this partition was assessed by repeating the clustering on 1,000 bootstrap resamples of the participants, and the large majority recovered the same two-metastate split. For each participant, the discrete state sequence was collapsed to a metastate sequence, from which the four dynamic functional network metrics were computed at the metastate level.

### 2.9 Energetic landscape and temporal irreversibility

A pairwise maximum-entropy model was fit to the binarized regional time series of each participant, which yields a coupling matrix (J) that defines the energetic landscape of the dynamics (Ezaki et al., 2017). The coupling structure was compared between groups by correlating the mean coupling matrices of the MS and HC groups. To quantify the directionality of the dynamics, temporal irreversibility was measured. Temporal irreversibility, also called the arrow of time, is the degree to which the sequence of activity differs from the same sequence played in reverse (Lynn et al., 2021). Irreversibility was computed for each connection and for the network as a whole, and groups were compared with a rank-based permutation test.

### 2.10 Statistical analysis

#### Group comparisons

For each of the 12 states, group differences in the four dynamic functional network metrics were assessed with a two-sided rank-based permutation test (10,000 permutations), the permutation analogue of the Mann-Whitney U test, which avoids distributional assumptions and is robust to the right-skewed lifetime and interval metrics. The same permutation test was applied to the subject-specific, dual-estimation state means and covariances to test for regional and connectivity differences between groups. Edge-level differences in state functional connectivity were localized with the network-based statistic (Zalesky et al., 2010). Within each state, a Welch t-statistic was computed for each of the 153 unique edges. Edges exceeding |t| = 3.0 were grouped into connected components, and component size was compared against a null distribution of the largest component from 5,000 group-label permutations. The four metastate-level metrics were compared with the same permutation test. As sensitivity analyses, the fractional-occupancy comparisons were repeated using occupancy from the continuous state posteriors rather than the most-likely-state sequence, and were re-tested in linear and logistic regression models adjusting for age and sex. A two-sided p-value less than 0.05 was considered to indicate statistical significance.

#### Clinical correlations

Within the MS group, associations between the per-state dynamic functional network metrics and clinical and cognitive variables (EDSS score, disease duration, processing speed, executive function, verbal memory, working memory, verbal fluency, and global cognition) were quantified with the partial Pearson correlation coefficient (r). Correlations with clinical variables were adjusted for age and sex, and correlations with cognitive variables for age, sex, and years of education.

## 3 RESULTS

### 3.1 Participant characteristics

The two groups were well matched (p > 0.05) for age (HCs, 36.2 ± 9.5 years, range 24-56; MS, 36.3 ± 8.7 years, range 20-50) and sex (8 men and 12 women per group). In the MS group, the median Expanded Disability Status Scale (EDSS) score was 3.5 (range 1.0-8.0), and the mean disease duration was 7.85 ± 5.31 years (range 3.0-22.0). Baseline demographic, clinical, and cognitive characteristics of the two groups are summarized in Table 1.

**TABLE 1.**
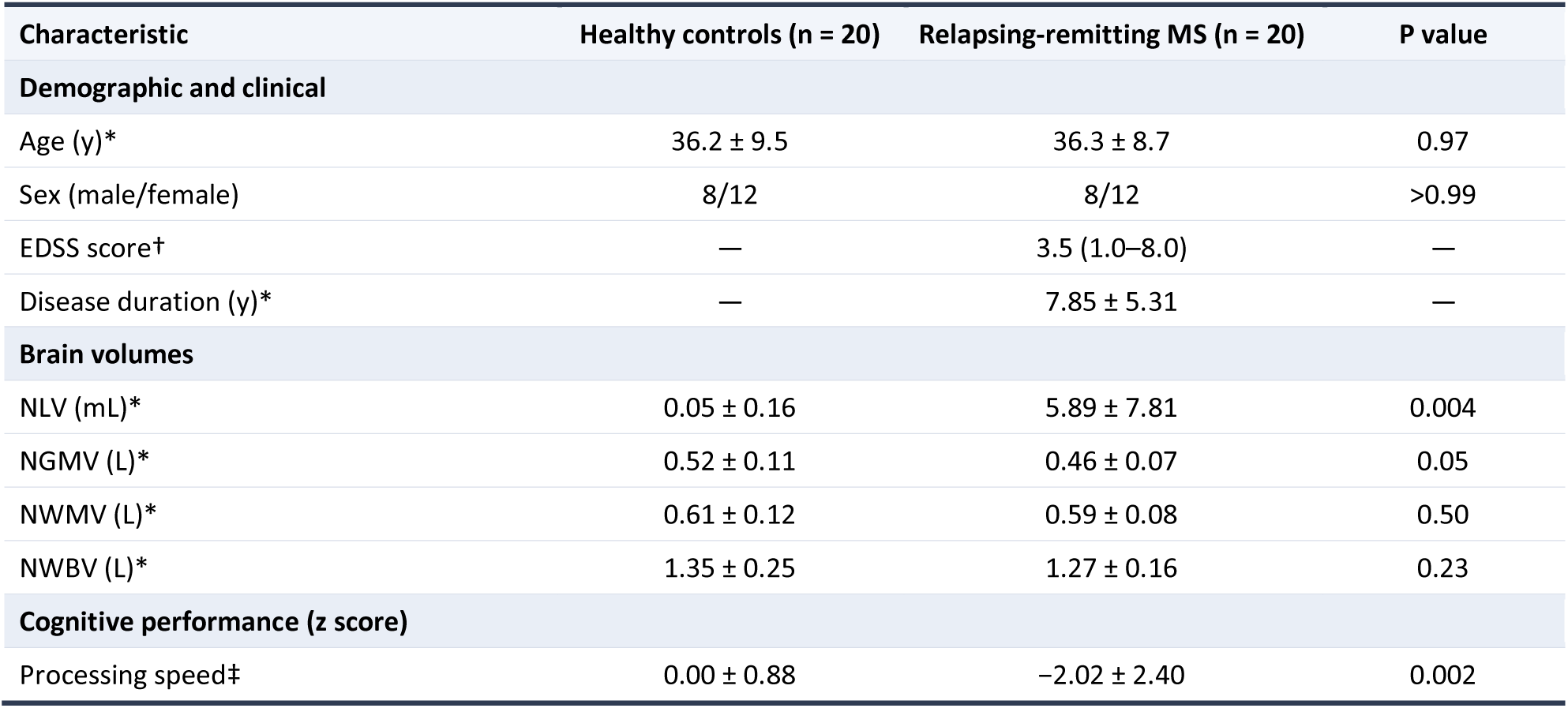
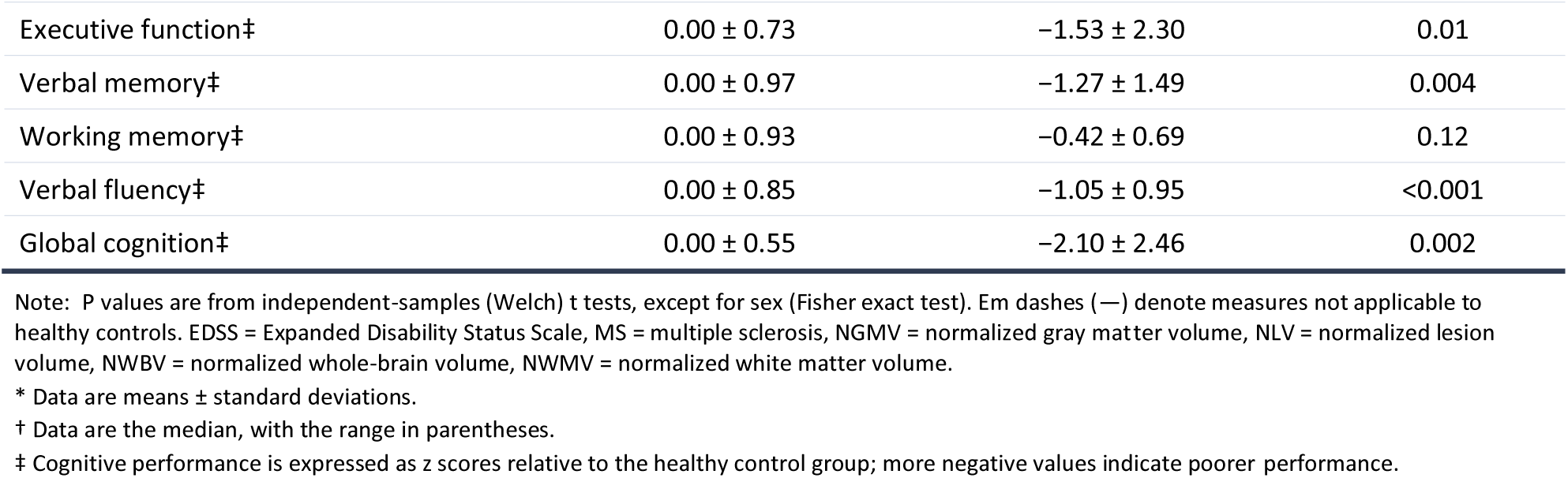
Characteristics and clinical status of study participants.

### 3.2 Characterization of brain states

The 12-state HMM produced a set of recurring whole-network activation states that people with MS and HC shared (Figure 2, A and B). Across the occupied states, the two groups showed similar mean activation maps, and the difference map was near zero in most region-by-state cells (Figure 2C). The few cells that differed were in subcortical and sensorimotor regions. Each state is a pattern across all 18 regions, shown in full in Figure 2. States are referred to by number and described by the region with the strongest activation, or by two regions where the two strongest were comparable. The description refers to the activation profile only. States 7 and 10 peaked on the postcentral gyrus (+0.70 and +0.51); in state 10 the left and right postcentral values were within 0.01 of each other, so this state is described by the structure rather than by one hemisphere. State 1 peaked on the right middle frontal gyrus (+0.92) and the right anterior cingulate cortex (+0.72), which were comparable, and is described by both. State 2 peaked on the left superior temporal gyrus (+1.08) and state 11 on the right middle frontal gyrus (+0.62). In state 3, activity fell below the network average in 16 of the 18 regions; the reduction was greatest in the cortex.

**FIGURE 1.**
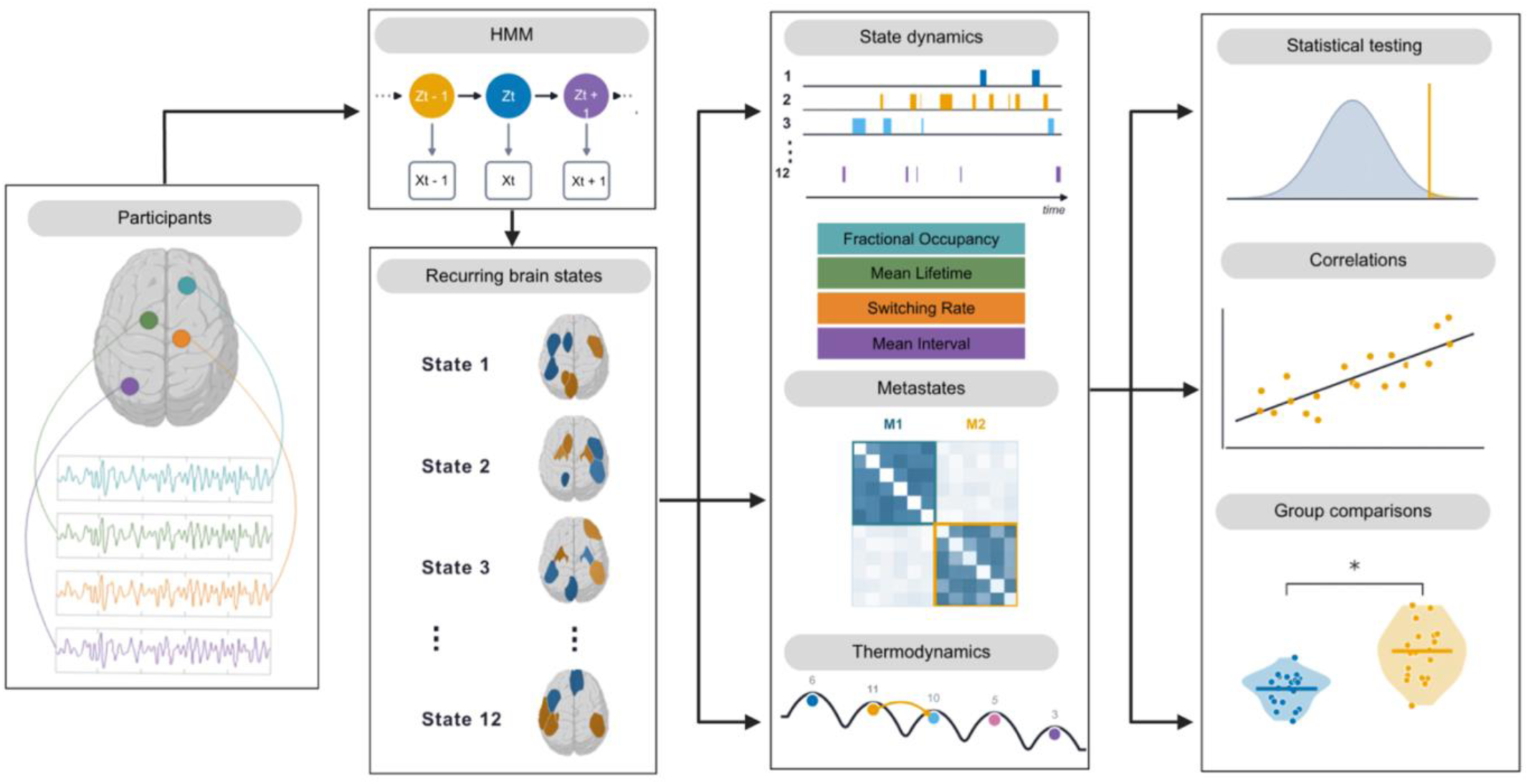
Overview of the analysis pipeline. Resting-state blood oxygen level-dependent (BOLD) time series were extracted from the 18 regions of the atrophy-based functional network in 20 people with MS and 20 healthy controls and preprocessed. A hidden Markov model was fitted to the group-concatenated data. The inferred state sequence yields the dynamic functional network metrics (fractional occupancy, mean lifetime, mean interval, switching rate, and transition probabilities), the grouping of states into two metastates, and the energetic-landscape measures. Group differences were assessed and state metrics were related to clinical scores within the MS group.

**FIGURE 2.**
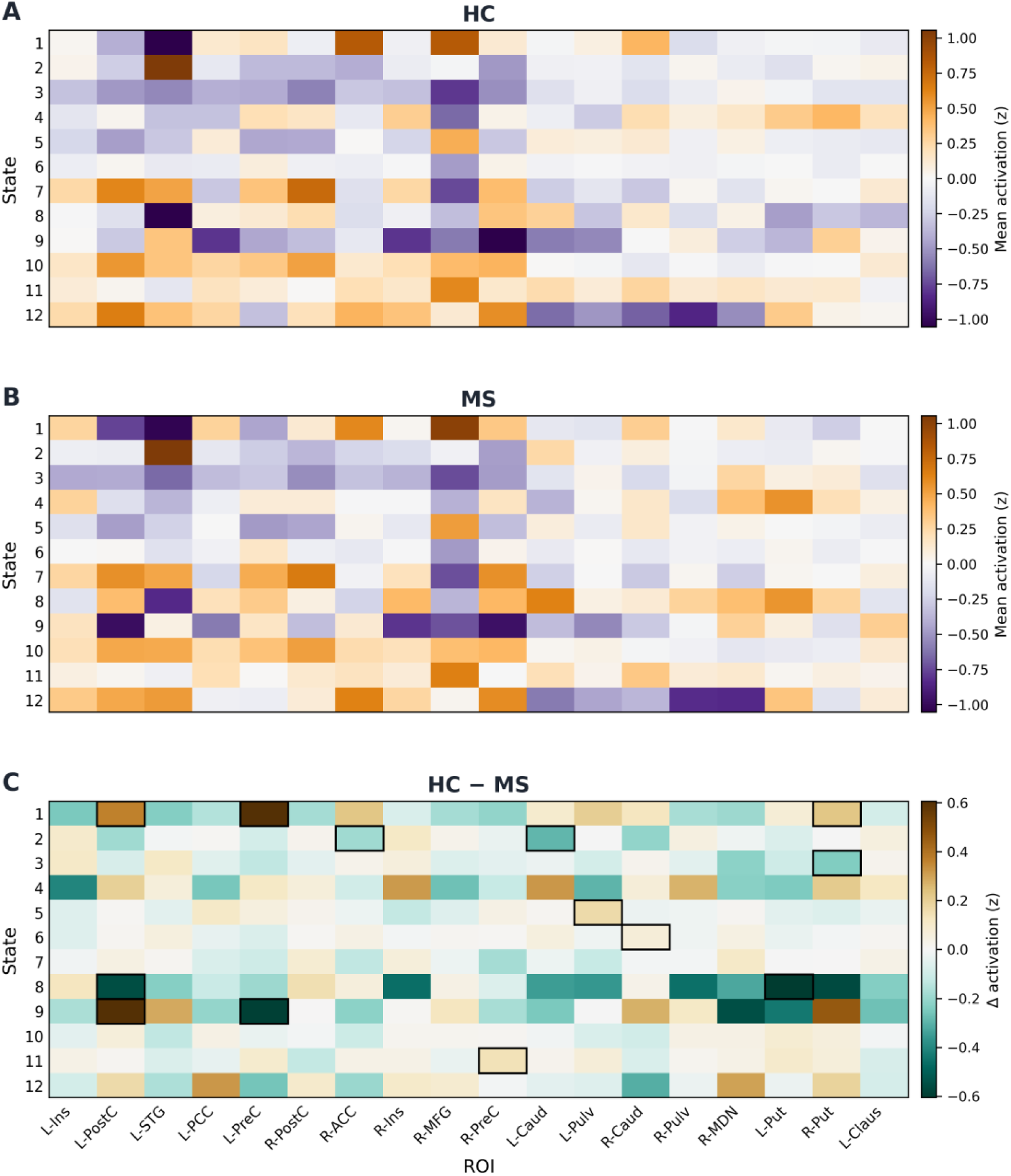
Mean activation of the HMM states in people with MS and HC. Heat maps show the mean activation (z-score) of each of the 12 states across the 18 ROIs in **(A)** healthy controls and **(B) people with multiple sclerosis (MS)**, and **(C)** the difference between groups (HC minus MS), in which orange/brown cells indicate greater activation in healthy controls, purple cells greater activation in MS, and black outlines mark the cells that differed significantly between groups (p < 0.05). The activation profiles were similar between groups, and the few differences were in subcortical and sensorimotor regions. ROI abbreviations: Ins = insula; PostC = postcentral gyrus; STG = superior temporal gyrus; PCC = posterior cingulate cortex; PreC = precentral gyrus; ACC = anterior cingulate cortex; MFG = middle frontal gyrus; Caud = caudate; Pulv = pulvinar; MDN = mediodorsal nucleus; Put = putamen; Claus = claustrum; L = left; R = right.

### 3.3 Altered temporal dynamics in multiple sclerosis

Between-group differences in the dynamic functional network metrics localized to four states (Figure 3; all states are shown in Figure S2). People with MS showed higher occupancy and longer lifetimes in state 3 (occupancy 0.088 in HC vs 0.130 in MS, p = 0.001; lifetime 8.3 vs 10.6 s, p = 0.02) and state 10 (occupancy 0.126 vs 0.165, p = 0.04; lifetime 5.9 vs 7.4 s, p = 0.04). In the opposite direction, people with MS showed lower occupancy of state 1 (0.045 vs 0.024, p = 0.007) and a lower switching rate into state 1 (p = 0.03). People with MS also visited state 7 less often, with a longer interval between visits (81.9 vs 91.2 s, p = 0.04). The overall rate of switching between states did not differ between groups.

**FIGURE 3.**
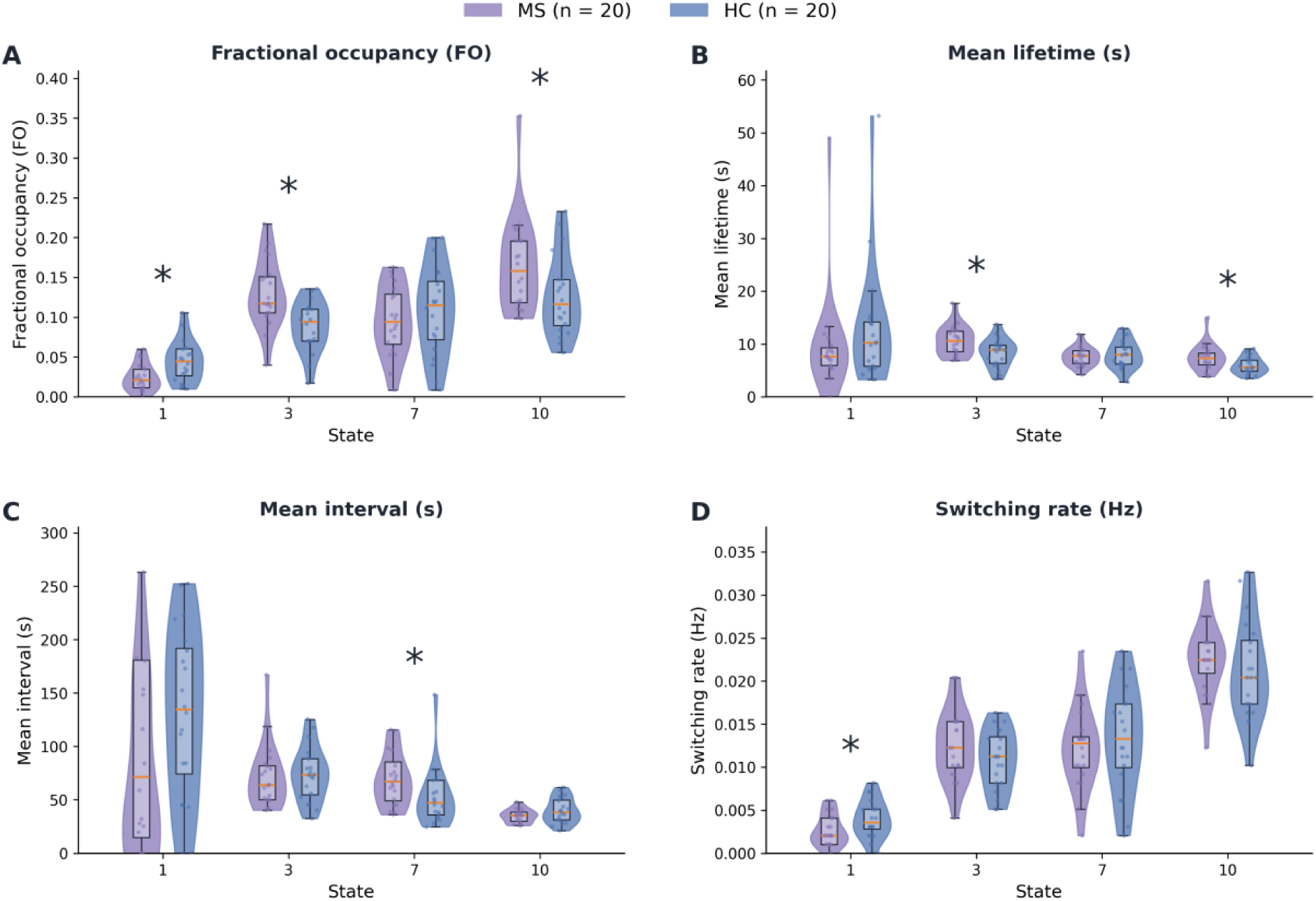
Dynamic functional network metrics of the states that differed between groups. Violin plots show the four metrics (fractional occupancy, mean lifetime, mean interval, and switching rate) for states 1, 3, 7, and 10 in people with MS (purple) and HC (blue). Boxes show the interquartile range and median. Asterisks mark a between-group difference (p < 0.05).

### 3.4 Associations with clinical and cognitive measures

Within the MS group, the per-state dynamic functional network metrics were correlated with the clinical and cognitive measures (Figure 4; partial correlation, n = 20). A longer interval between visits to state 3 was associated with greater disability (r = 0.57; p = 0.01), and a higher switching rate in state 10 with poorer verbal fluency (r = −0.50; p = 0.04). A longer lifetime of state 11, which peaked on the right middle frontal gyrus, was associated with greater disability (r = 0.54; p = 0.02).

**FIGURE 4.**
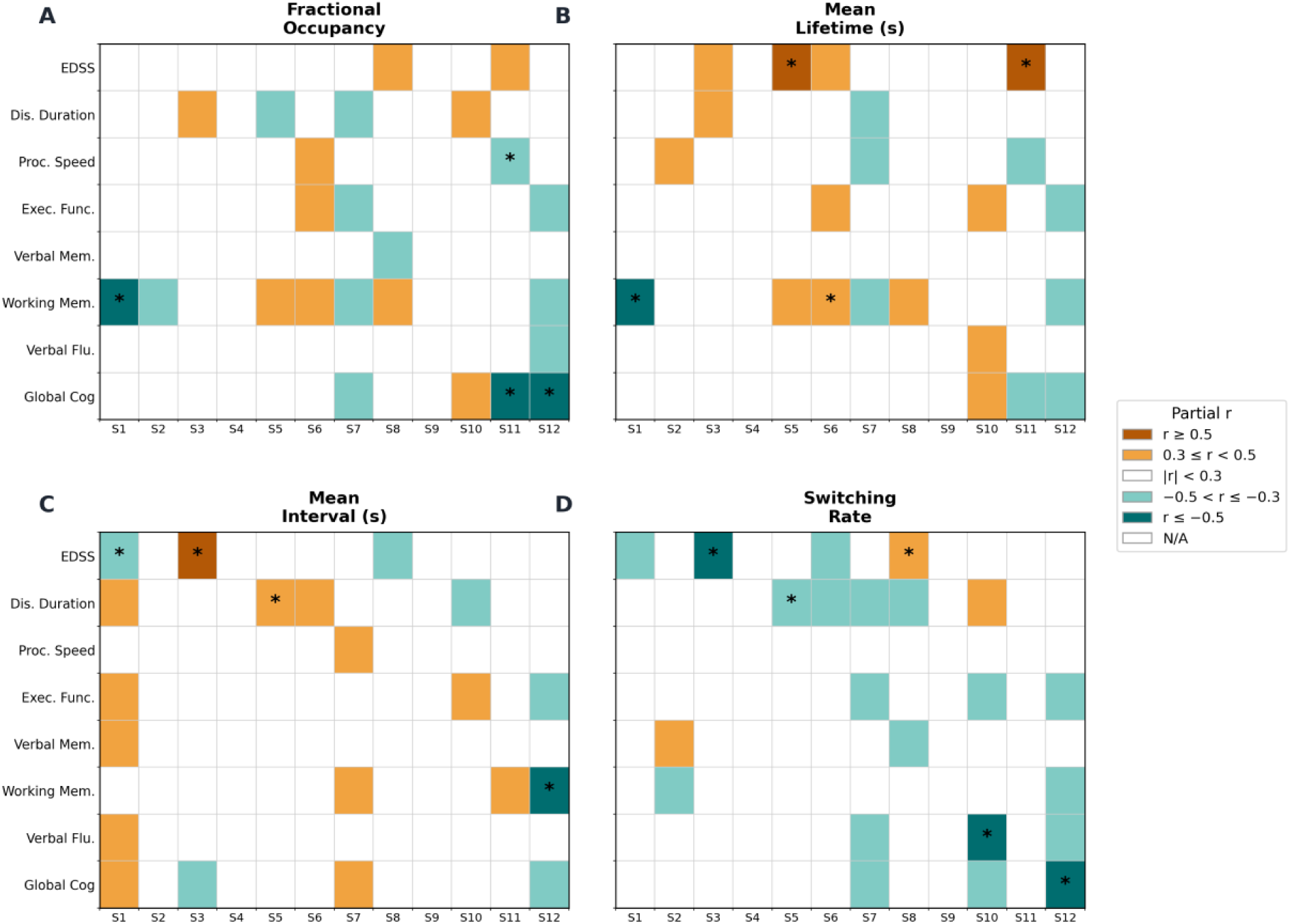
Within-MS correlations between dynamic functional network metrics and clinical and cognitive measures. Heat maps show the partial correlation within the MS group (n = 20) between the four per-state dynamic functional network metrics (fractional occupancy, mean lifetime, mean interval, and switching rate) and the clinical and cognitive variables, adjusted for age and sex (clinical variables) or age, sex, and education (cognitive variables). Cell color indicates the strength and direction of each correlation, and asterisks mark p < 0.05. EDSS = Expanded Disability Status Scale.

### 3.5 Metastate organization

The active states were grouped into two metastates by clustering the transition probability matrix (Figure 5, A-C). One metastate (M1) comprised states 2, 3, 6, 7, 8, and 10, and activated the postcentral and precentral gyri (+0.10 and +0.07) while suppressing the right middle frontal gyrus (−0.33). The other (M2) comprised states 1, 5, and 11, and activated the right middle frontal gyrus and anterior cingulate cortex (+0.67 and +0.31) while suppressing the left superior temporal gyrus (−0.50). Only fractional occupancy differed between groups (Figure 5D). People with MS spent a greater fraction of time in M1 (0.67 vs 0.64 in HC) and a smaller fraction in M2 (0.33 vs 0.36; p = 0.04). Mean lifetime, mean interval, and switching rate did not differ between groups (Figure 5, E-G).

**FIGURE 5.**
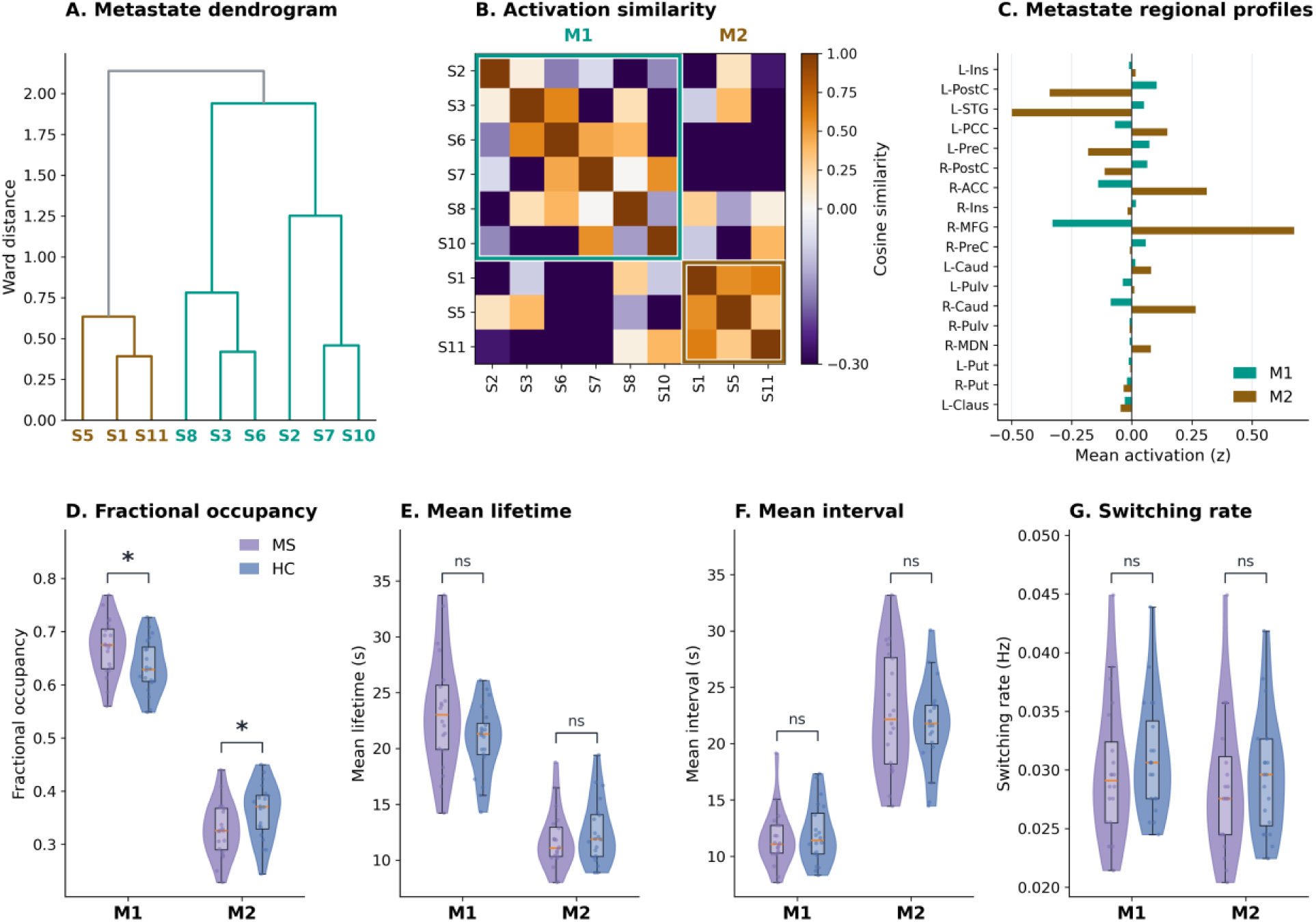
Metastate organization of the states. **(A)** Ward dendrogram of the state mean-activation profiles, with leaves and branches colored (teal: M1, brown: M2) by the metastate to which each state was assigned by clustering the transition probability matrix. **(B)** Cosine-similarity matrix of the state activation profiles ordered by metastate, with colored boxes delineating the two metastates. **(C)** Mean activation of each metastate across the 18 AFN regions. **(D-G)** Fractional occupancy, mean lifetime, mean interval, and switching rate for each metastate in people with multiple sclerosis (purple) and healthy controls (blue); fractional occupancy differed between groups (asterisk in D), whereas the other three metrics did not. ROI abbreviations: Ins = insula; PostC = postcentral gyrus; STG = superior temporal gyrus; PCC = posterior cingulate cortex; PreC = precentral gyrus; ACC = anterior cingulate cortex; MFG = middle frontal gyrus; Caud = caudate; Pulv = pulvinar; MDN = mediodorsal nucleus; Put = putamen; Claus = claustrum; L = left; R = right.

### 3.6 Energetic landscape and temporal irreversibility

The energetic organization of the dynamics was largely preserved in MS. The pairwise coupling between regions that defines the energetic landscape was highly similar between the two groups, with a strong correlation between the group mean coupling matrices (r = 0.90; Figure 6A). Temporal irreversibility quantifies how distinguishable the observed sequence of activity is from the same sequence played in reverse. A strongly irreversible process runs in a clear direction in time, whereas a reversible one looks the same forwards and backwards. Temporal irreversibility was lower in the MS group than in controls (Figure 6B), so the dynamics were less directed in time in MS. This reduction was concentrated in subcortical connections (Figure 6C). The largest reductions were between the caudate and putamen, and between the striatum or pulvinar and the cingulate and insular cortex.

**FIGURE 6.**
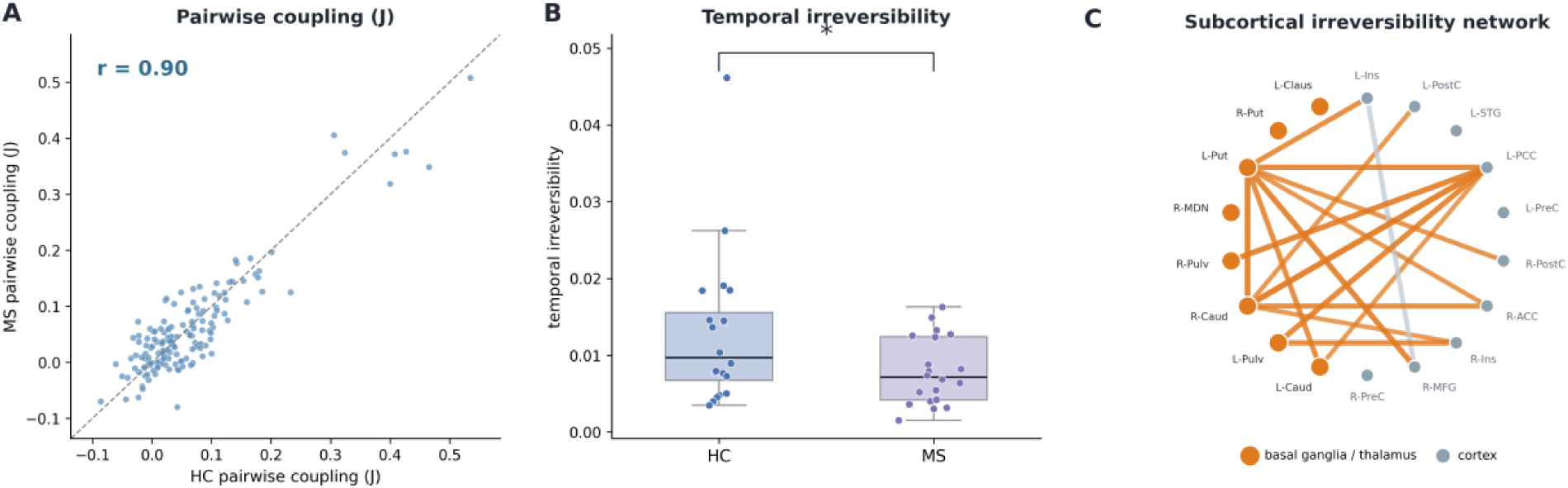
Energy-based analyses of the dynamics. **(A)** Pairwise coupling (J) between regions in the energetic landscape in people with multiple sclerosis (MS) plotted against healthy controls (HC); the coupling structure was correlated between groups (r = 0.90), consistent with a similar landscape. **(B)** Temporal irreversibility, or arrow-of-time asymmetry, of the network in each participant, with boxes showing the median and interquartile range; irreversibility was lower in MS (p = 0.048). **(C)** The connections with the largest between-group reduction in irreversibility (healthy controls minus MS), shown on the network; the strongest links (orange) all involved basal-ganglia or thalamic nodes, consistent with a loss concentrated in subcortical circuits. ROI abbreviations: Ins = insula; PostC = postcentral gyrus; STG = superior temporal gyrus; PCC = posterior cingulate cortex; PreC = precentral gyrus; ACC = anterior cingulate cortex; MFG = middle frontal gyrus; Caud = caudate; Pulv = pulvinar; MDN = mediodorsal nucleus; Put = putamen; Claus = claustrum; L = left; R = right. The median and interquartile range are shown in (B) because the control distribution included a single outlying value; the lower irreversibility in MS does not depend on this value.

## 4 DISCUSSION

We characterized functional network dynamics involving atrophy-prone regions in MS by applying hidden Markov modeling via the AFN model. Within this framework, people with MS drew on the same recurring states as healthy controls but distributed time across those states differently. Occupancy and mean lifetime were higher in states 3 and 10, and occupancy was lower in state 1, which peaked on the right middle frontal and anterior cingulate cortex. Time was therefore redistributed among the states, while the overall rate of switching did not differ between groups. The same redistribution appeared at the metastate level. Within the MS group, dynamic functional network metrics were associated with physical disability and cognitive impairment. In addition, the energetic landscape was similar between groups while temporal irreversibility of subcortical connections was lower in MS. This pattern suggests reduced directed drive within circuits linking the basal ganglia and thalamus. These findings are in line with models of MS as a disorder of distributed network disruption rather than of focal lesion burden alone. The findings also suggest that dynamic functional connectivity has the potential to capture clinically relevant network alterations in MS beyond static functional connectivity. Assessment of functional network dynamics may therefore provide further insights into development of network-based imaging approaches for monitoring disease in MS.

The group differences lay in the timing of the states rather than in the regions those states engaged. Functional connectivity studies in MS have reported widespread but heterogeneous abnormalities that vary in direction and location across cohorts (Hidalgo de la Cruz et al., 2021; Tahedl et al., 2018). In this sample the spatial patterns were largely the same in both groups, differing only in a few subcortical and sensorimotor regions (Figure 2C). In contrast, the dynamic functional network metrics differed between groups in four states (Figure 3). Functional connectivity averaged over an entire acquisition cannot distinguish a change in the spatial pattern of a state from a change in the time spent in that state (Hutchison et al., 2013). Prior dynamic functional connectivity studies in MS have accordingly reported differences in state dwell time and in the frequency of state transitions (Broeders et al., 2022; d’Ambrosio et al., 2020; von Schwanenflug et al., 2023). The present findings suggest that, within the AFN, alterations in MS are expressed mainly in the timing of the states.

The states that differed in timing between groups peaked on cortical regions of the AFN. States 7 and 10 peaked on the postcentral gyrus, and state 1 peaked on the right middle frontal gyrus and anterior cingulate cortex. State 3 also differed in timing but has no regional peak. State 8 peaked on the left caudate but did not differ in timing between groups. The striatum and thalamus instead carried the reduction in temporal irreversibility (Figure 6C). Altered timing and altered directionality were therefore observed in different parts of the network.

The changes in network dynamics took the form of prolonged dwelling in two states. People with MS showed higher fractional occupancy and longer mean lifetimes in states 3 and 10, so that each visit to either state lasted longer. The overall switching rate did not differ between groups. That extra time came from other states: occupancy of state 1 was lower, and state 7 was visited less often. The group difference therefore lay in the distribution of time across states rather than in the overall rate of switching. Less frequent switching and a smaller distance travelled through the state repertoire were reported in cognitively impaired than in cognitively preserved patients (d’Ambrosio et al., 2020); more frequent network reconfiguration in impaired patients was reported in a second cohort (Broeders et al., 2022). Both studies relate switching to cognitive status within the patient group. This relation is in line with our within-patient correlations between the dynamic functional network metrics and disability and cognition. Greater persistence of connectivity states with increasing disease severity, from healthy controls to relapsing-remitting and progressive MS, has also been reported with a sliding-window, clustering-based Markov model (Tessadori et al., 2021).

The largest increase in fractional occupancy in MS was observed in state 3. Cortical activity fell below the network average, most markedly in the right middle frontal gyrus, the pre- and postcentral gyri, and the left superior temporal gyrus. Thalamic and striatal activity remained near baseline. The additional time spent in state 3 may therefore reflect more frequent or prolonged periods of reduced cortical engagement rather than sustained recruitment of any single region (d’Ambrosio et al., 2020). In the MS group the metrics of state 3 related to physical disability rather than to processing speed, even though processing speed is among the domains most consistently affected in MS (Chiaravalloti & DeLuca, 2008).

Within the MS group, the dynamic functional network metrics that correlated with the clinical measures belonged to states 3, 10, and 11. A longer interval between visits to state 3 was associated with greater disability (r = 0.57), and a higher switching rate into state 10 with poorer verbal fluency (r = −0.50). Both describe less sustained engagement of these states in participants with worse clinical status, consistent with reports linking worse clinical outcome in MS to states entered less often and held less steadily (Broeders et al., 2022; d’Ambrosio et al., 2020; von Schwanenflug et al., 2023). A longer mean lifetime of state 11, which peaked on the right middle frontal gyrus, was also associated with greater disability (r = 0.54). Taken together, these associations suggest that the dynamic functional network metrics capture variation in clinical status within the MS group.

At a coarser scale, the states organized into two metastates (Figure 5). M1 comprised states 2, 3, 6, 7, 8, and 10, which activated the postcentral and precentral gyri. M2 comprised states 1, 5, and 11, which activated the right middle frontal and anterior cingulate cortex. People with MS spent more time in M1 and less in M2. In healthy participants the time spent in each metastate is a stable, heritable trait related to cognition (Vidaurre et al., 2017). The decrease in time spent in M2 is consistent with the reduced cortical engagement reported in patients with MS and cognitive decline (Broeders et al., 2022; d’Ambrosio et al., 2020). At both the state and the metastate level, therefore, the MS group spent less time engaging right middle frontal and anterior cingulate cortex than controls did.

The activation differences and the loss of irreversibility both converged on subcortical circuits. The few activation differences between groups were in subcortical and sensorimotor regions (Figure 2C), and the loss of temporal irreversibility was concentrated in subcortical connections, both within the striatum and between subcortical nodes and the cingulate and insular cortex (Figure 6). The arrow of time of neural activity reflects the directional, hierarchical organization of information flow, so a weaker arrow of time points to a flatter and less directed hierarchy (de la Fuente et al., 2023; Lynn et al., 2021). Irreversibility was reduced in nodes of the cortico-striato-thalamo-cortical loops, which are largely closed and unidirectional (Alexander et al., 1986). A reduction in irreversibility within these loops is consistent with a flattening of the directed drive that normally propagates activity around the loops. A similar pattern has been reported in Parkinson’s disease, a disorder centered on the basal ganglia and thalamus, in which brain dynamics shift toward equilibrium and the hierarchy of cortico-subcortical loops becomes flatter (Sadeghi et al., 2025). Our results are consistent with a similar pattern in MS and may relate to the structural vulnerability of the striatum and thalamus, which show early gray-matter atrophy, iron deposition, and disconnection in the disease (Schoonheim et al., 2022).

Monitoring of disease progression in MS relies largely on white matter lesion burden, which correlates only weakly with clinical disability (Barkhof, 2002). Measures of network dynamics are obtained from a single resting-state acquisition without a task and could in principle be acquired alongside standard-of-care imaging. The dynamic functional network metrics reported here differed between groups at the level of individual states and were associated with disability and cognition within the MS group. These associations support further evaluation of the dynamic functional network metrics as candidate markers.

Limitations of this study include a cross-sectional design, which prevents inference about disease progression, and a small single-site sample of participants with relapsing-remitting disease and generally low disability. Age and sex are associated with functional connectivity (Ferreira & Busatto, 2013; Ritchie et al., 2018), and age, sex, and education are associated with cognitive test performance in MS (Parmenter et al., 2010). These variables may therefore confound associations in a sample of this size. The within-group correlations were therefore adjusted for age and sex, and the correlations with cognitive measures were additionally adjusted for education. Replication in a larger, multi-site cohort with out-of-sample testing is needed to establish whether the state-level and subcortical effects can serve as individual-level markers. Resting-state dynamic measures can also partly reflect sampling variability, head motion, and arousal rather than genuine state changes (Laumann et al., 2017; Lurie et al., 2020). To mitigate this, we applied denoising and nuisance regression, and the fractional-occupancy differences persisted after adjustment for age and sex. Relating these measures and the loss of subcortical irreversibility to regional atrophy, lesion load, and structural disconnection would further tie the functional findings to the underlying anatomy, as would computing transition energies from each participant’s structural connectome.

## 5 CONCLUSION

Dynamic functional alterations involving a distributed network of atrophy-prone regions in MS were associated with disease worsening. We identified changes in the timing of the network states rather than in the regions those states engaged, while the overall rate of switching did not differ from healthy controls. Within the MS group, state timing was associated with physical disability and cognitive impairment. Altered timing was observed in cortical states, whereas altered directionality was observed in subcortical connections. This pattern is consistent with altered network dynamics rather than altered network architecture. Network-based dynamic measures may provide a basis for quantitative, noninvasive monitoring of neurodegeneration in MS.

## Data Availability Statement

The primary structural and functional imaging data can be made available via a Data Transfer and Use Agreement with The University of Texas Health Science Center at San Antonio upon reasonable request (Contact: Peter T. Fox).

## SUPPLEMENTARY MATERIAL

**FIGURE S1.**
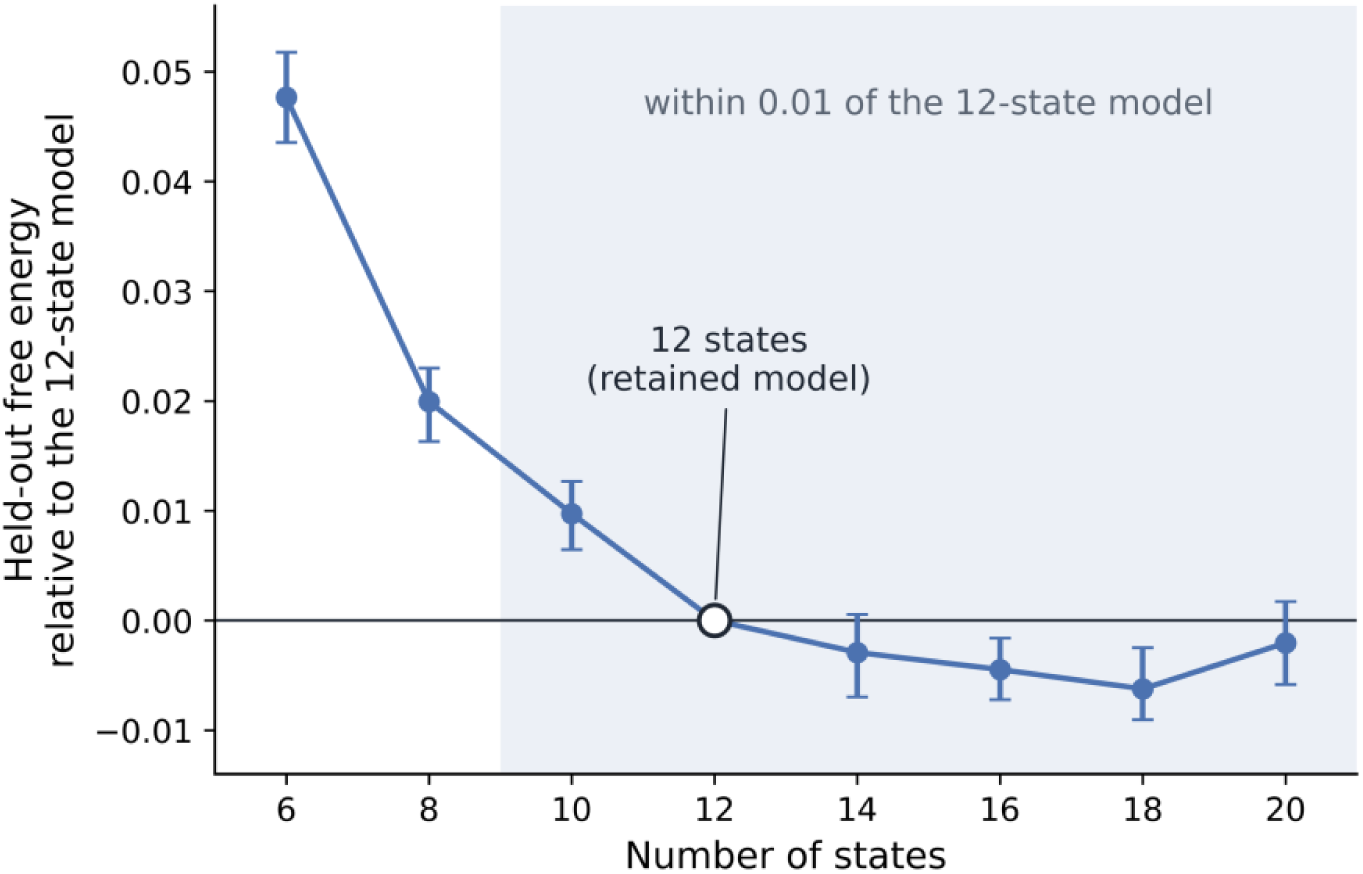
Selection of the number of HMM states. Free energy in the held-out validation sample for models with 6 to 20 states, shown relative to the 12-state model (open circle). Points show the median difference across 800 training configurations, and error bars show the bootstrap 95% confidence interval. Lower values indicate better fit. Free energy decreased from 6 to 10 states, and models with 10 to 20 states (shaded) differed from the 12-state model by 0.01 or less.

**FIGURE S2.**
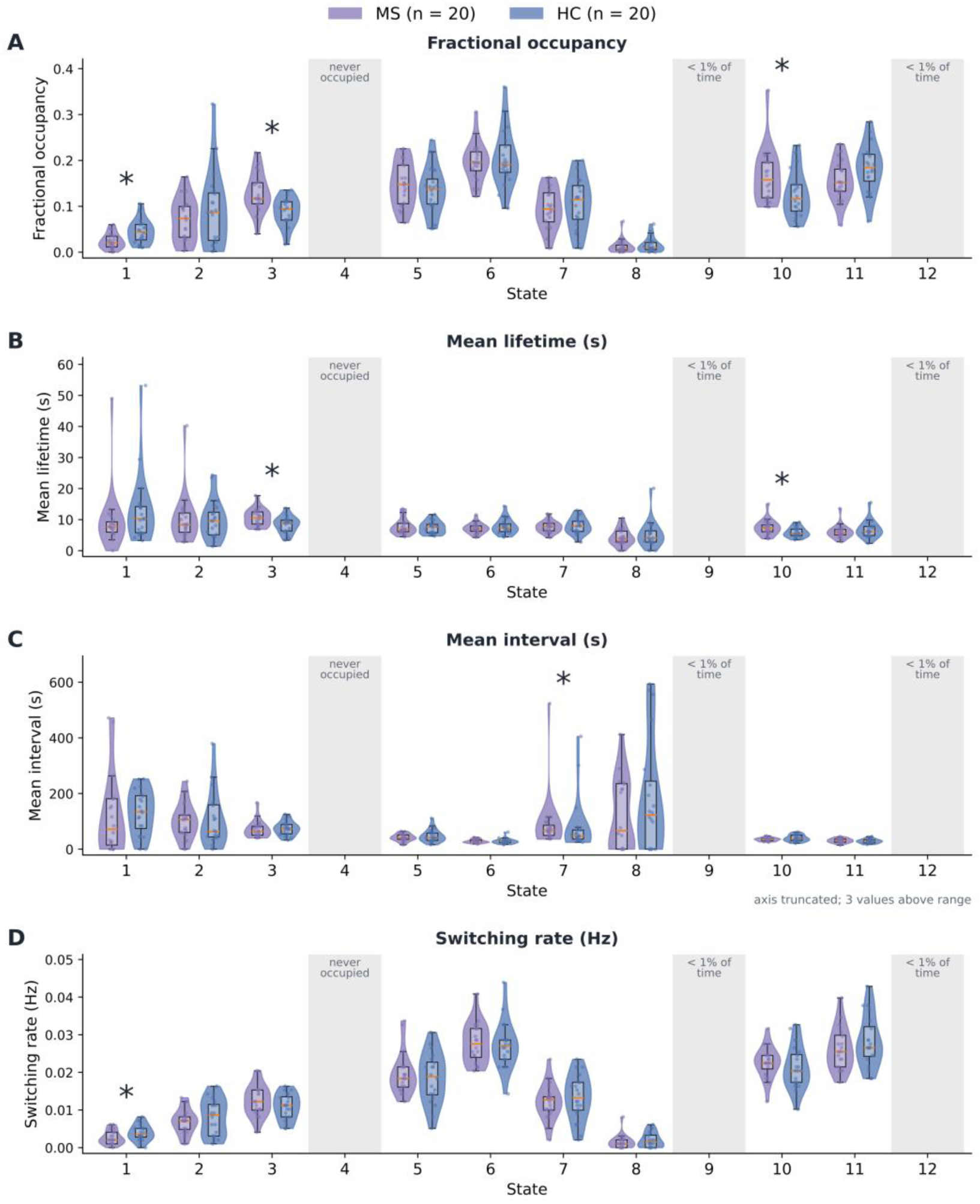
Dynamic functional network metrics of all 12 states. Violin plots show (A) fractional occupancy, (B) mean lifetime, (C) mean interval, and (D) switching rate for each state in people with MS (purple) and HC (blue). Boxes show the interquartile range and median. Asterisks mark a between-group difference (p < 0.05, rank-based permutation test). Gray bands mark the states that were not tested: state 4 was never occupied, and states 9 and 12 were each occupied for less than 1% of time points.

**FIGURE S3.**
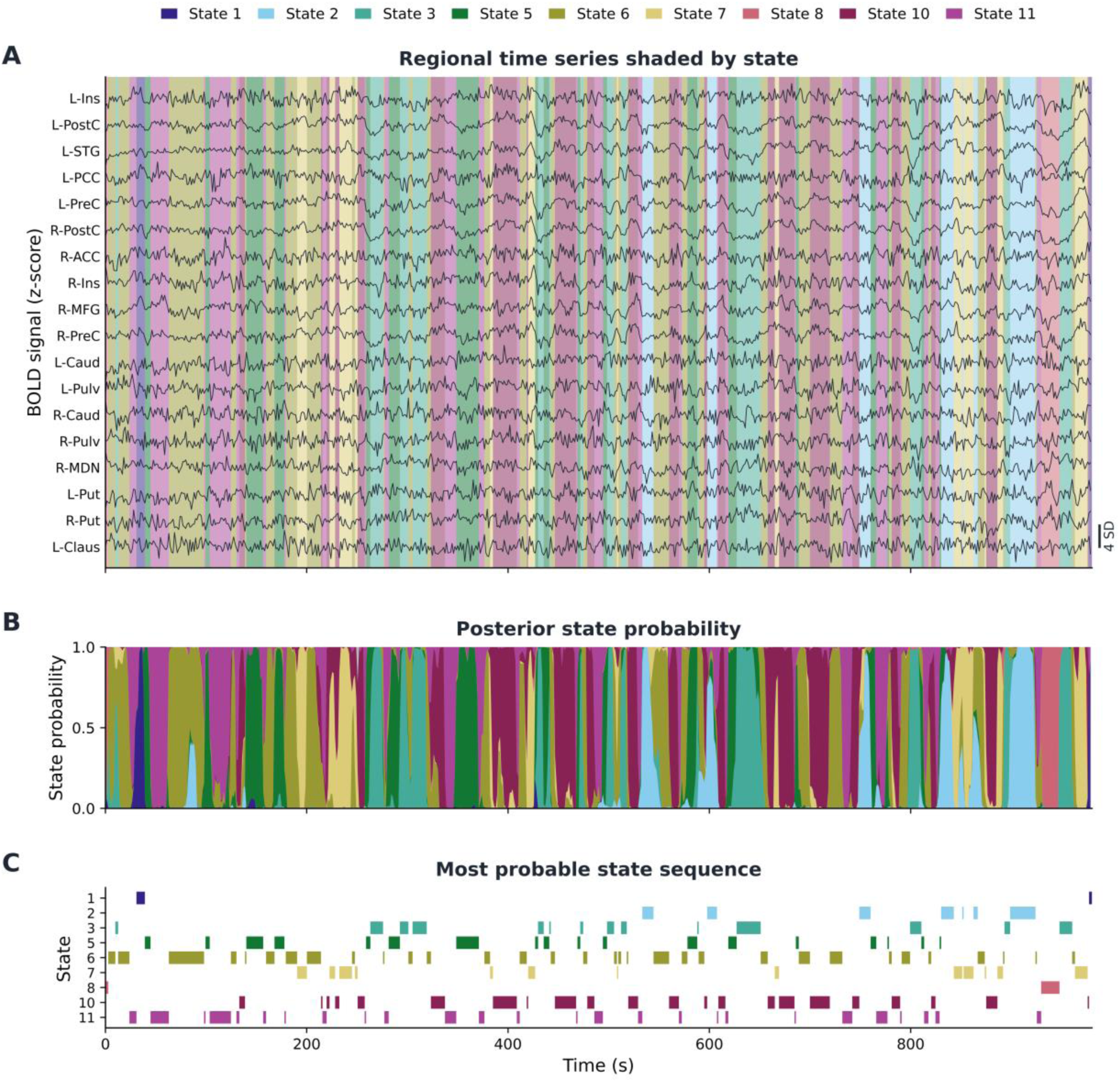
Regional time series and inferred state sequence in one participant with MS. (A) Standardized BOLD time series of the 18 ROIs over the 980-second acquisition (700 volumes); background color indicates the state assigned to each time point. Nine of the 12 states appear because this participant did not enter states 4, 9, or 12. (B) Posterior probability of each state at each time point. (C) Most probable state sequence. The participant shown is the one whose fractional-occupancy profile was closest to the MS group mean. ROI abbreviations: Ins = insula; PostC = postcentral gyrus; STG = superior temporal gyrus; PCC = posterior cingulate cortex; PreC = precentral gyrus; ACC = anterior cingulate cortex; MFG = middle frontal gyrus; Caud = caudate; Pulv = pulvinar; MDN = mediodorsal nucleus; Put = putamen; Claus = claustrum; L = left; R = right.

